# The superior colliculus and the steering of saccades toward a moving visual target

**DOI:** 10.1101/133736

**Authors:** Laurent Goffart, Aaron Cecala, Neeraj Gandhi

## Abstract

Following the suggestion that a command encoding the expected here-and-now target location feeds the oculomotor system during interceptive saccades, we tested whether this command originates in the deep superior colliculus (SC). Monkeys generated saccades to targets that were static or moving along the preferred axis, away from (outward) or toward a fixated target (inward) with a constant speed (20°/s). Vertical and horizontal motions were also tested. Extracellular activity of 57 saccade-related neurons was recorded in 3 monkeys. The movement field (MF) parameters (boundaries, center and firing rate) were estimated after spline fitting the relation between the saccade amplitude and the average firing rate of the motor burst. During radial motion, the inner MF boundary shifted in the same direction as the target motion for some neurons, not all. During vertical motion, both lower and upper boundaries were shifted upward during upward motion whereas the upper boundary only shifted during downward motions. For horizontal motions, the medial boundaries were not changed. The MF center was shifted only for outward motion. Regardless of the motion direction, the average firing rate was consistently reduced during interceptive saccades. Our study shows an involvement of the saccade-related burst of SC neurons in steering the gaze toward a moving target. When observed, the shifts of MF boundary in the direction of target motion correspond to commands related to antecedent target locations. The absence of shift in the opposite direction shows that SC activity does not issue predictive commands related to the future target location.

**SIGNIFICANCE STATEMENT:** By comparing the movement field (MF) of saccade-related neurons between saccades toward static and moving targets, we show that the motor burst issued by neurons in the superior colliculus does not convey commands related to the future location of a moving target. During interceptive saccades, the active population consists of a continuum of neurons, ranging from cells exhibiting a shift in the center or boundary of their MF to cells which exhibit no change. The shifts correspond to residual activity related to the fact that the active population does not change as fast as the target in the visual field. By contrast, the absence of shift indicates commands related to the current target location, as if it were static.

## INTRODUCTION

The primate oculomotor system has been used as a model to understand the neuronal processes underlying the localization of an object in the external world and the generation of movements toward its location (Goffart, 2017). In most studies, the stimulus was static, leaving unexplored the processes generating saccades toward a moving target. An involvement of the deep superior colliculus (SC) and caudal fastigial nucleus (CFN) is however suggested by the emission of bursts of action potentials by their neurons during interceptive and catch-up saccades aimed at a moving target (Keller et al., 1996; Fuchs et al., 1994). Moreover, their anatomical situation between the cerebral and cerebellar cortices where neurons responsive to the motion of a target are found (Cassanello et al., 2008; Robinson and Fuchs, 2001) and the saccade-related premotor neurons in the reticular formation (Scudder et al., 2002; Sparks, 2002) corroborates their involvement.

According to the "dual drive" hypothesis, interceptive saccades are driven by a combination of commands issued by these two subcortical structures (Optican 2009). The locus of SC activity encodes the location where the target first appears whereas the CFN component encodes the command related to the target motion after the collicular "snapshot" (see also Optican & Pretegiani 2017). This hypothesis rests upon the observation that the centers of the movement field (MF) of SC neurons (i.e., the amplitude and direction of saccades associated with the most vigorous burst) shifts to larger amplitudes during saccades toward a target moving away from the central visual field (Keller et al., 1996). However, the magnitude of the shift spans over a notable range since some neurons exhibit no change (see their Fig. 3A). This scattering suggests instead that the population of neurons which burst during interceptive saccades consists of a continuum of cells ranging from neurons issuing commands related to past locations of the target (cells with a shift) to neurons issuing commands related to its current location (cells with no shift). Thus, as an alternative to the dual drive hypothesis, the "remapping" hypothesis proposed that the population of active neurons in the SC does not correspond to a snapshot, but expands across the SC (Fleuriet et al. 2011). In other words, the supplementary command envisioned by the dual drive hypothesis would be incorporated within the SC itself, making its output a possible origin of the expected here-and-now command that has been proposed to feed the saccade premotor system during interceptive saccades (Fleuriet & Goffart 2012).

**Figure 1:**
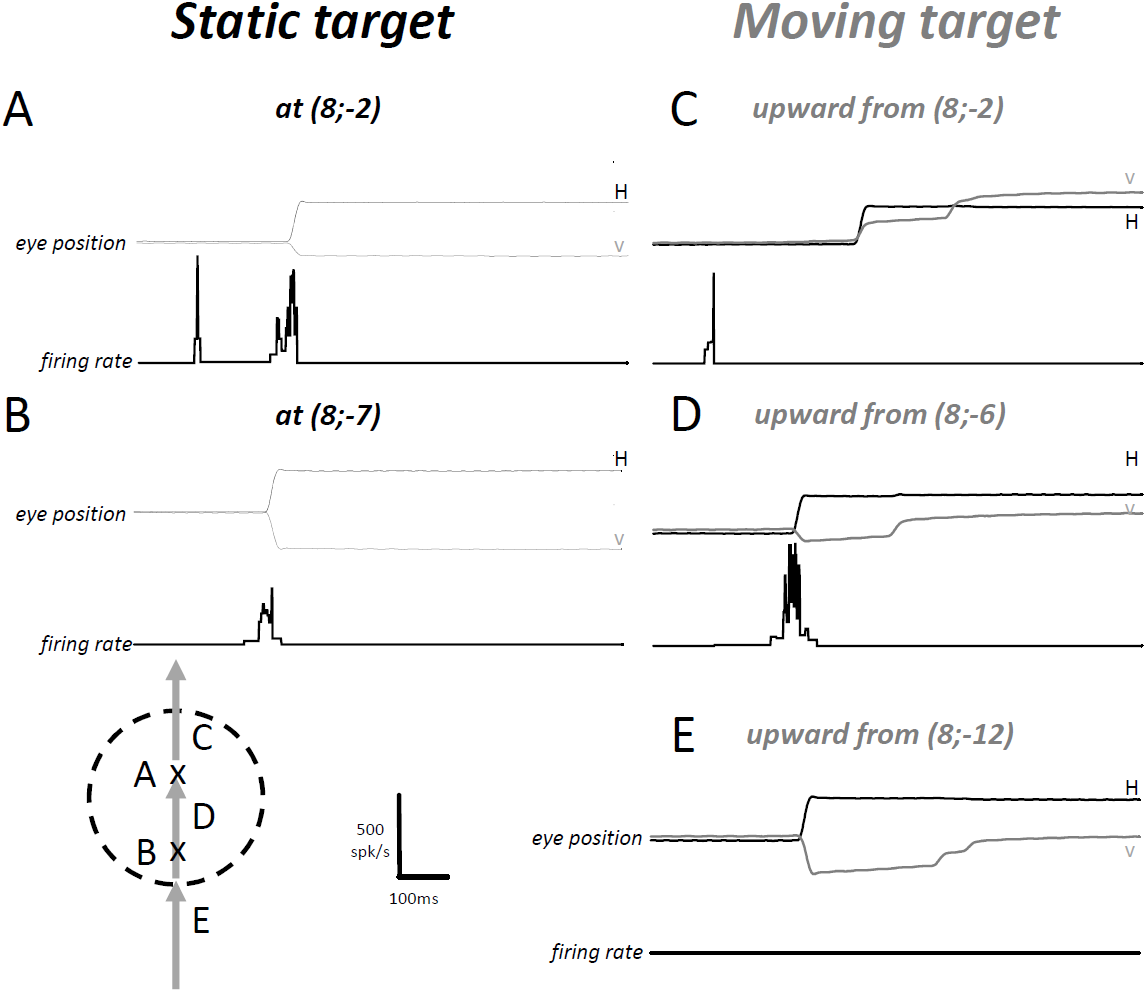
Illustration of the firing rate of a SC visuomotor neuron during single trials. A-B: Visual and saccade-related activity following the appearance of a static target at different locations (Cartesian coordinates) of the right visual field. C-E: Activity of the same neuron when the target moves upward at the same horizontal eccentricity. In A and C, the target appears at a location corresponding to the center of the neuron’s movement field (MF). In D, the saccade is aimed at the same location as in A: the visual response is absent because the moving target appears outside the neuron’s response field. In E, the saccade is aimed at the same location as in B: the neuron does not fire when the target moves.

**Figure 2:**
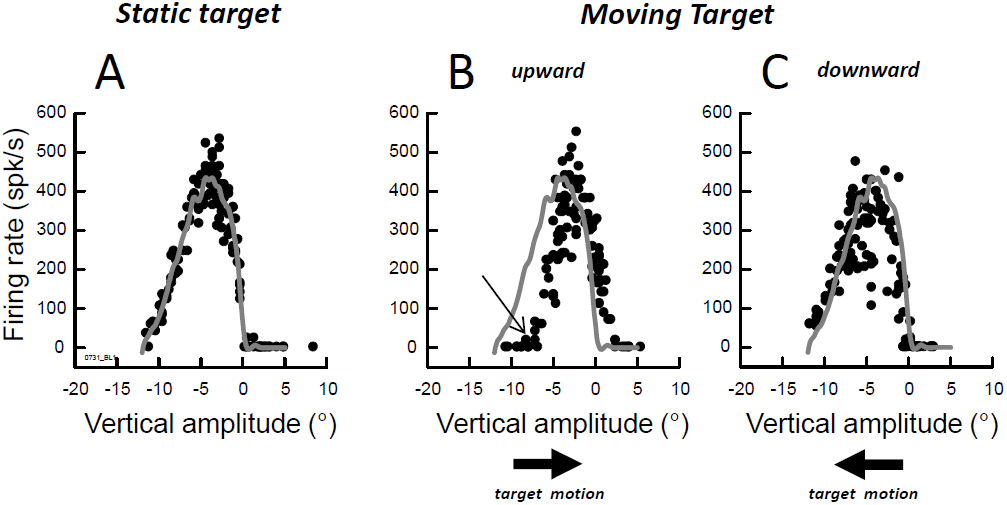
Movement field of the same neuron as in Figure 1 during saccades toward targets located on an axis parallel to the vertical meridian. A: static target; B: target moving upward; C: target moving downward. The arrow in B shows the shift in the lower boundary of the MF.

**Figure 3:**
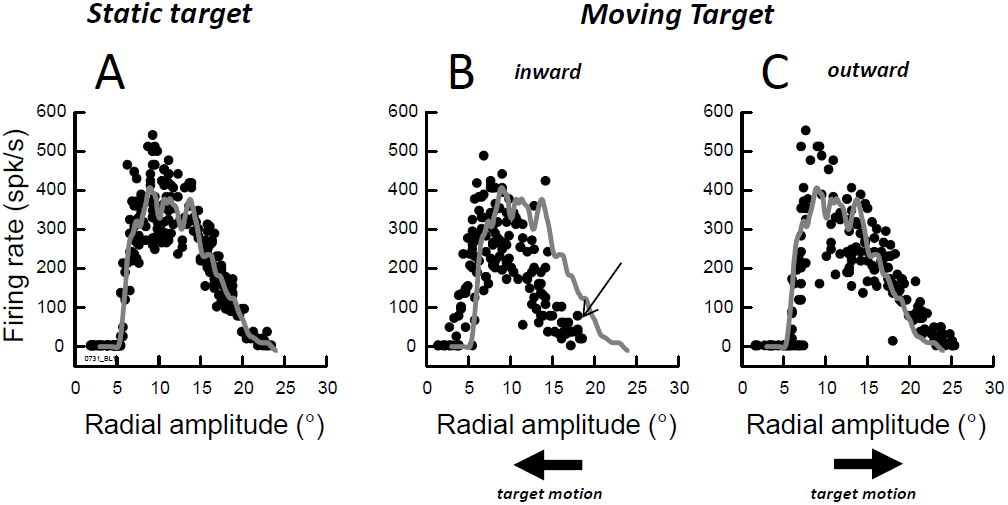
Movement field of the same neuron as in Figures 1 and 2 during saccades toward targets located along the radial axis of its MF. A: static target; B: target moving inward (toward the fixation target); C: target moving outward (away from the fixation target). The arrow in B shows the shift in the outer boundary of the MF when the target moves toward it.

The goal of this study was to evaluate these hypotheses. We also examined whether SC neurons bursting during interceptive saccades issue commands related to future locations of the target along its motion path, i.e., locations which are going to be reached. Such a possibility would be indicated by shifts in the *boundaries* of the MF in the direction *opposite* to the target motion, an option which was not addressed in the study of Keller et al. (1996) since they focused on the MF centers. Furthermore, we complemented the electrophysiological characterization of saccade-related neurons in the SC by comparing their MF between saccades to static and moving targets. Our results show a continuum of neurons, ranging from cells which exhibit a shift in the boundary (or center) of their MF to cells which do not exhibit any change. When shifts were observed, they were always in the same direction as the target motion, never in the opposite direction. This absence of shift in the opposite direction indicates no recruitment of neurons which issue commands related to any future target location. When they are observed, the shifts correspond to a residual activity due to the fact that the locus of active neurons across the SC does not change as fast as the target in the visual field. The observation of cells with no shift is consistent with their involvement in steering the saccade toward the current location of a moving target, as if it were static.

## MATERIALS AND METHODS

### Subjects and surgical procedures

All surgical and experimental protocols were approved by the University of Pittsburgh Animal Care and Use Committee and performed in accordance with the National Institutes of Health Guide for the Care and Use of Laboratory Animals. Three adult rhesus monkeys (*Macaca mulatta*; Male: BB & BL; Female: WI) underwent aseptic surgeries to secure a small head-restraint device to the skull, cement a stainless steel chamber over a craniotomy, and attach a Teflon-coated stainless steel wire (search coil) on the sclera of one eye. The chamber was placed stereotaxically on the skull, slanted posteriorly at an angle of 38° in the sagittal plane. This approach allowed access to both SC and permitted electrode penetrations roughly perpendicular to the SC surface. Antibiotics and analgesics were administered postoperatively as detailed in an approved protocol.

### Behavioral tasks and experimental apparatus

After full recovery, the subjects were trained to sit in a primate chair with their head restrained and a sipper tube placed near the mouth for reward delivery. They were subsequently trained to perform standard oculomotor tasks involving stationary targets. The monkeys were not previously trained to pursue moving targets, which were introduced only during the recording sessions. Visual stimuli, behavioral control, and data acquisition were implemented by a custom-built program that uses LabVIEW conventions on a real-time operating system supported by National Instruments (Austin, TX) (Bryant and Gandhi, 2005). Each animal sat inside a frame containing two alternating magnetic fields that induced voltages in the search coil thereby permitting measurement of horizontal and vertical eye orientations (Robinson 1963). Visual targets were red dots subtending ∼0.5° of visual angle that were displayed on a 55 inch, 120 Hz resolution LED monitor.

Every trial began with the illumination of an initial target (T0) that the subjects were required to fixate for a variable duration (300-700ms, 100ms increments). Trials were aborted if the gaze direction deviated beyond a computer-defined window (3° radius) surrounding T0. If fixation was maintained, then T0 was extinguished and another target (T1) was simultaneously presented in the visual periphery. During static trials, the subjects were rewarded for orienting their gaze within a window that surrounded T1 with a radius of 3-6° for a minimum of 350 ms. During motion trials, target T1 moved at a constant speed of 20°/sec immediately after it appeared on the screen. The reward window associated with T1 was elliptical with a long axis that extended from the starting position of T1 to at least 5° beyond its final position. The subjects were required to be within this window for at least 500ms before receiving reward. The starting position and the direction of target motion depended upon the movement field properties of the recorded cell as determined during static trials (see below).

### Single-unit recording and movement fields

Tungsten microelectrodes (Microprobe) were used to record extracellular activity from the intermediate and deep layers of SC. The SC was identified online by the presence of distinctive bursts of activity associated with flashes of room lights and saccades as well as identifiable saccade-related cells during static trials. After we isolated a single saccade-related neuron, we estimated the boundaries of its movement field by pseudo-randomly presenting targets and observing peak firing rates displayed online by the acquisition software. Once the optimal vector was approximated, a series of static target locations was chosen along either 1) an imaginary line that passed through the center of the movement field and the initial target T0; or 2) an imaginary line that passed through the center of the movement field and parallel to the vertical meridian; or 3) an imaginary line that passed through the center of the movement field and parallel to the horizontal meridian. Approximately 75-100 static trials were collected before static and motion trials were pseudo-randomly intermixed. The starting positions of moving targets (T1 ini) were located along the same axis used for the targets during static trials. Target motion could be radial (inward or outward relative to T0), vertical (upward or downward relative to T1 ini) or horizontal (rightward or leftward relative to T1 ini). Data were collected across the three axes in block mode. Introducing variability in the location of T1 ini during the motion trials, as well as the natural variability in the subjects’ reaction times, allowed the collection of neural data during interceptive saccades that fell both within and outside of the boundaries of the movement field as defined during static trials.

### Data set and analysis

The horizontal and vertical eye positions for each trial were digitized and stored with a resolution of 1 ms and then analyzed off-line analysis with a custom software and Matlab. The onset and offset of saccades were identified using a velocity criterion of 15°/s. Saccade metrics (amplitude, peak velocity, latency, etc.) reported here were obtained by measuring the first saccade (primary saccade) made after the presentation of T1 (equivalently, offset of T0). The primary saccade needed to occur between 100ms and 500ms after the offset of T0 in order to be considered for further analysis.

The present study concerns the discharge properties of 57 neurons which fired a burst of action potentials during saccades. Response fields were obtained by plotting firing rate (calculated as the number of spikes per second during a period beginning 20 ms before saccade onset and continuing until 10 ms before saccade end) as a function of horizontal, vertical, or radial saccade amplitude during either the static or motion trials. The boundaries and optimal vector encoded by the MF were estimated from a smoothing spline fit of the data with the curve-fitting toolbox in Matlab. For each neuron, the same spline parameter was used for fitting the data of both tasks. The boundary was defined as the saccade amplitude from which the neuron starts firing with a rate larger than 30 spikes/s. When the saccade-related burst was preceded by a prelude activity, the threshold was adjusted to the minimal value that characterizes the burst onset. The Wilcoxon test (P<0.05) was used to test for statistically significant differences in MF properties across neurons between the saccades toward a static and a moving target.

## RESULTS

Figure 1A illustrates the firing rate of a typical visuomotor SC neuron during a static target trial. The first phasic response occurred approximately 100 ms after the onset of the visual target and was followed by a second, more vigorous burst timed with the saccade toward its location. The neuron also produced a weaker burst during saccades whose amplitude and direction slightly deviated from the neuron’s preferred vector (Fig. 1B); the visual response was absent for this particular location. In response to a target moving upward at the same horizontal eccentricity, the neuron’s discharge was different. When the target motion started from the location that elicited vigorous visual and perisaccadic responses during the static condition (Fig. 1A), the visual response was not followed by the saccade-related burst (Fig. 1C). Thus, the response of this neuron could signal the presence of the target within its response field, but it did not participate to the population activity which drives this particular interceptive saccade. The cell was active during saccades whose vectors matched the vectors that elicited the most vigorous perisaccadic bursts with a static target (compare Fig. 1A to Fig. 1D). Another observation is the absence of firing when the monkey made an interceptive saccade whose vector was associated with a perisaccadic burst if the target had been static (compare Fig. 1B to Fig. 1E). During this particular condition, the neuron was silent even though the saccade vector belonged to the movement field measured with static targets (hereafter referred to as “static MF”) and even though the target was going to enter within this MF.

Figure 2 plots a slice through the MF of the same cell during three target conditions: static (2A), moving upward (2B) and downward (2C). The three MFs were generated by presenting targets along a vertical axis situated at a horizontal eccentricity of 8° to the right. During these target conditions, saccades had horizontal amplitudes ranging from 7.2 to 9.2°. The center of the static MF was identified during rightward saccades with a small (-4.4°) downward component (Fig. 2A); the discharge of this neuron was weaker when the saccade deviated from this preferred vertical amplitude. Estimated by a spline fitting procedure, the lower and upper boundaries of the static MF as were −11.2° and 0.1°, respectively. Compared to the static MF, the center and the boundaries of the dynamic MF (Fig. 2B) were shifted upward (toward positive values) during saccades made to a target moving upward (center: −2.3°, shift Δ = 2.1°; lower boundary = −7.3°, Δ = 2.9°; upper boundary = 2.4°, Δ = 2.3°). When the target appeared below the lower edge of the static MF and moved upward toward the center of the MF, the neuron did not fire unless the interceptive saccade involved a vertical component larger than −7.3° (see arrow in Fig. 2B). Thus, instead of emitting spikes that would promote the foveation of a target which was going to enter in its MF, the neuron remained silent. Likewise, when the vertical amplitude of the interceptive saccade exceeded the amplitude corresponding to the upper boundary of the static MF (0.1°), instead of pausing and facilitating the generation of saccades with a larger upward component, this neuron emitted spikes, biasing the population of active neurons with a command encoding an oblique downward vector. While differences between the static and dynamic MFs were clearly visible during saccades directed to a target moving upward, changes were barely visible in the saccade-related burst of this neuron when the target moved downward (Fig. 2C). Thus, the effects of a moving target on the MF properties of this particular neuron were consistent with the "dual drive" hypothesis when the saccades were made to a target moving upward, and with the "remapping" hypothesis when they were made to a target moving downward.

Next, we examine the saccade-related burst of the same neuron during saccades made along the radial axis of its MF. During saccades to static targets, the neuron fired during saccades of radial amplitudes ranging from 5.5° (inner boundary) to 20.7° (outer boundary) with the most vigorous bursts occurring for 8.9° saccades (Fig. 3A). During saccades to a target moving from the peripheral to the central visual field (inward saccades), the entire MF was shifted toward smaller amplitude values (Fig. 3B). When the target started its motion from outside the MF and moved inward, the neuron did not fire unless the monkey made a 17° saccade (see arrow in Fig. 3B). Thus, instead of emitting spikes that would promote the reduction of saccade amplitudes, the neuron remained silent. Moreover, although no firing was observed during small saccades toward static targets with eccentricity less than 5°, the neuron discharged during small inward saccades. A small shift of the entire MF was also observed in the direction of the target motion during outward saccades: the outer boundary shifted toward larger amplitudes (Δ = 2.1°; Fig. 3C) whereas the inner boundary barely changed (Δ = 0.4°).

Many of the cells that we recorded exhibited open movement fields, so only the proximal boundary could be identified. Figure 4 shows four examples of such neurons where the movement field exhibited a shift in boundary (consistent with the "dual drive" hypothesis) whereas Figure 5 shows examples of neurons where the shift was absent or barely visible (consistent with the "remapping" hypothesis). Fig. 4 shows the movement fields during saccades made to a static target (black symbols) or to a target moving (grey symbols) along an axis orthogonal to the vertical meridian (A: rightward motion), a radial axis (B: outward motion) or an axis perpendicular to the horizontal meridian (C: downward motion; D: upward). For each of these neurons, the boundary of the MF is shifted in the same direction as the target motion. By contrast, Fig. 5 shows examples of neurons which exhibited no shift or barely visible shift in the MF boundary during interceptive saccades (like in Fig. 3C). Some of them exhibited a lower firing rate during saccades made to the center of the movement field (Fig. 5A-C and F). However, this reduced firing rate was not observed during small (Fig. 5A,D) or large (Fig. 5C,F) saccades.

**Figure 4:**
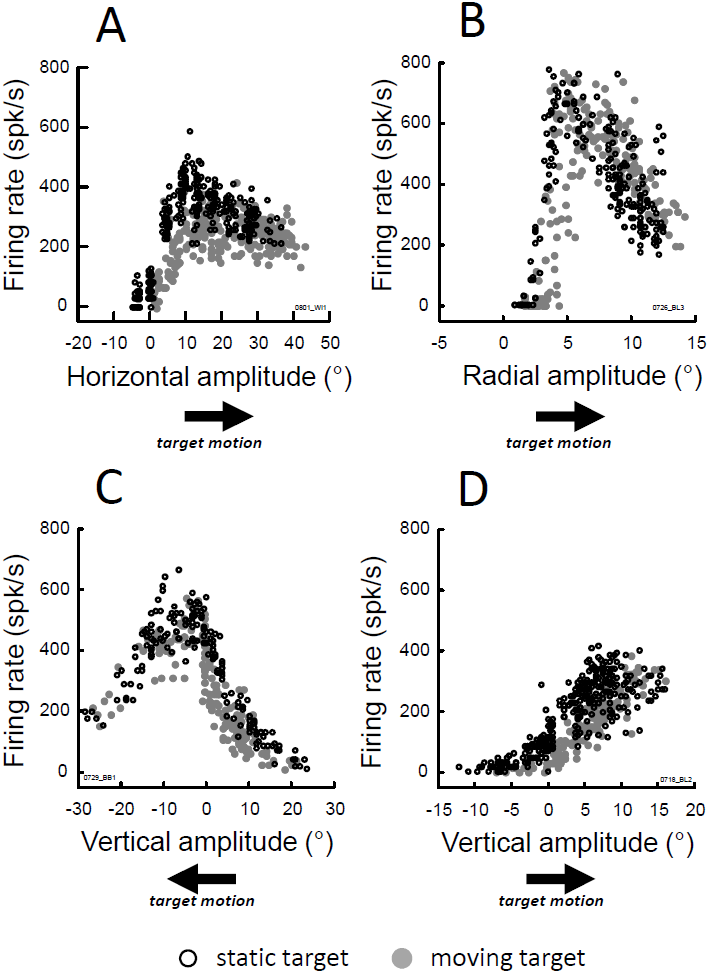
Movement fields of four other neurons exhibiting a shift during saccades toward a moving target (grey) in comparison to saccades toward a static target (black). A: target moves to the right; B: target moves outward along the radial axis; C: target moves downward; D: target moves upward.

**Figure 5:**
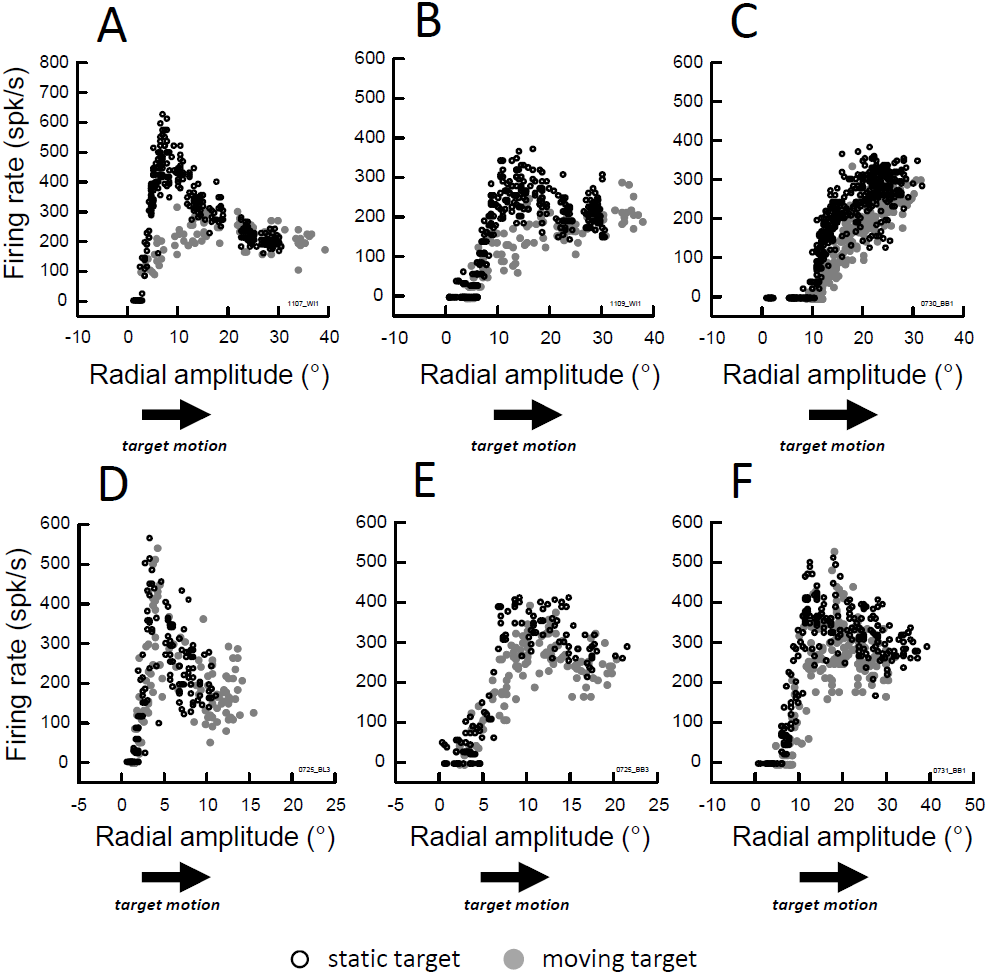
Movement fields of six other neurons exhibiting no shift, neither in the center nor the inner boundary. Grey: firing rate during interceptive saccades, black: firing rate during saccades toward a static target.

Figure 6 compares, for all neurons, the boundaries of static MF to those of MF measured during saccades made toward a target that moved radially (A), horizontally (B), or vertically (C: upward or D: downward) across their MF. In comparison to the static target conditions, the inner boundary shifted toward small amplitude values when the saccades were made to a target that moved inward, i.e., toward the central visual field (Fig. 6A, left graph; average difference=-1.6+/-1.6 deg, non-parametric Wilcoxon test, P<0.05). During outward motion (Fig. 6A, right graph), a small, but significant shift toward larger amplitude values, in the same direction as the target motion, was also observed (0.7+/-0.8 deg, P<0.05). When the target moved horizontally across the MF (Fig. 6B), no significant difference in the medial boundary were observed during leftward (0.9+/-2.2 deg; P-value = 0.25) or rightward (1.4+/-3.8 deg; P-value = 0.29) motion. The absence of significant difference is likely due to the small sample of neurons recorded during this motion condition of target motion. In contrast, when the target moved upward (Fig. 6C), a shift in the same direction as the target motion was observed for the lower boundary (leftmost graph in C; 2.3+/-2.0 deg, P<0.05). For the upper boundary (rightmost graph in C), the difference failed to reach our threshold of statistical significance (1.1+/-2.0 deg; P-value=0.07). During downward target motion (Fig. 6D), a significant shift was observed for the upper boundary (-1.9+/-1.7 deg, P<0.05; rightmost graph in D) but not for the lower boundary (0.0+/-1.2 deg, P-value=0.81; leftmost graph in D,). In summary, average shifts in the MF boundaries were observed but not in every condition. Crucially, whenever a significant difference was found between the static and dynamic MFs, the shift was always in the same direction as the target motion.

**Figure 6:**
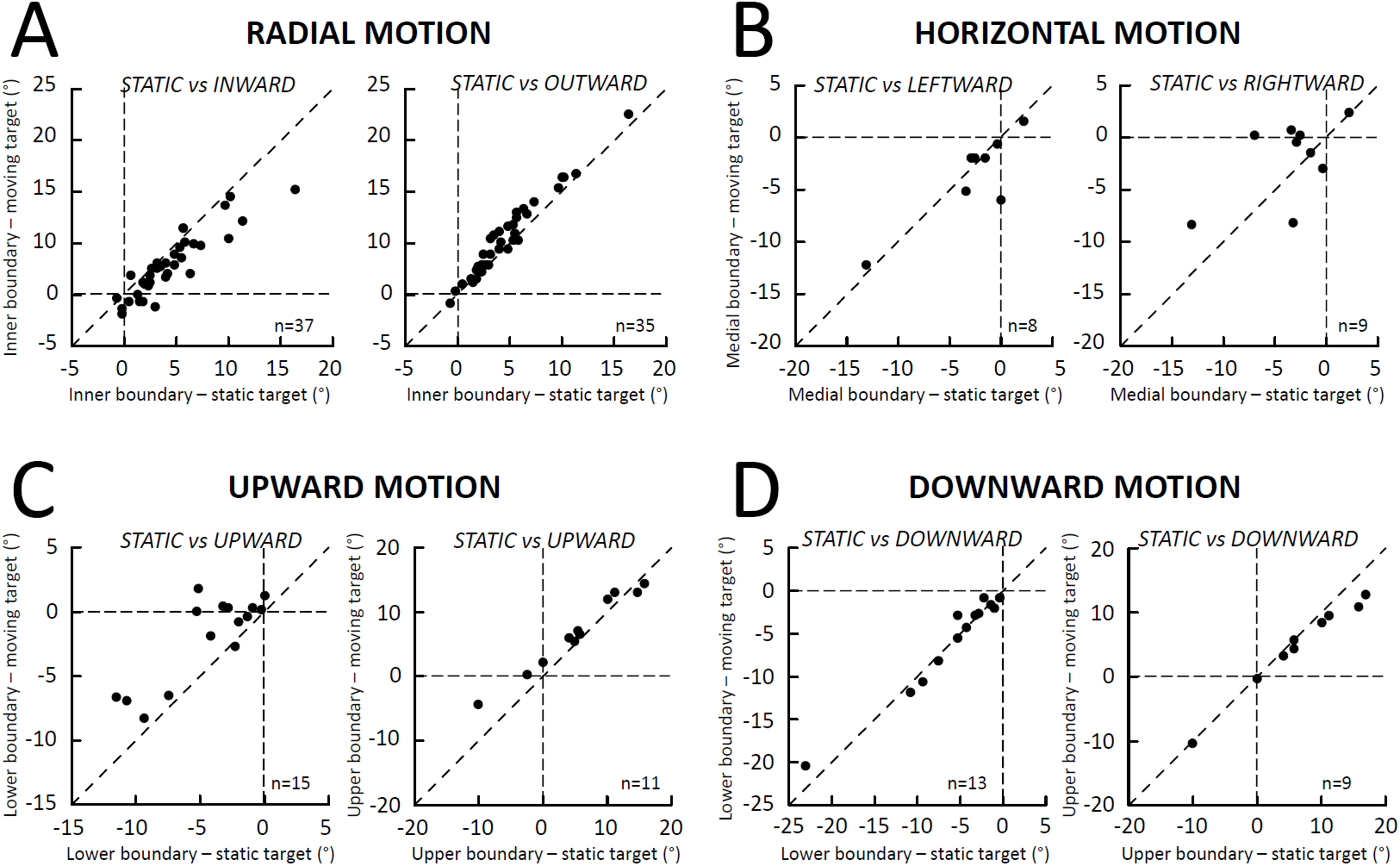
Comparison of the MF boundaries between saccades toward a static target (abscissa) and saccades toward a target (ordinate) moving along the radial axis (A), a horizontal axis (B) and a vertical axis passing through the MF center (C and D). The moving target moves upward in C, downward in D.

While Keller et al. (1996) did not describe the MF boundaries, they reported a shift in MF centers during saccades made toward stimuli moving outward; other directions of target motion were not tested. Figure 7 complements and extends their study by comparing the preferred amplitude values during radial (panel A), vertical (B) and horizontal (C) target motions. The center of MF significantly changed during outward saccades (Fig. 7A, right graph; average difference = 3.0+/-4.2 deg, P < 0.05). No consistent shift was observed during inward saccades (0.4+/-3.6 deg; P-value = 0.54). During vertical motions (Fig. 7B), a shift was observed when the target moved upward (3.1+/-3.2 deg, P < 0.05; rightmost graph) but not when it moved downward (-0.2+/-3.3 deg, P-value = 0.81; leftmost graph). During horizontal target motion (Fig. 7C), we could not detect any significant change for leftward (-3.6+/-4.4; P-value=0.052) and rightward (1.7+/-4.3 deg; P-value=0.29) motions. In summary, shifts in the MF center were observed but not in every condition. Whenever a significant difference was found between the static and dynamic MF, the shift was always in the same direction as the target motion.

**Figure 7:**
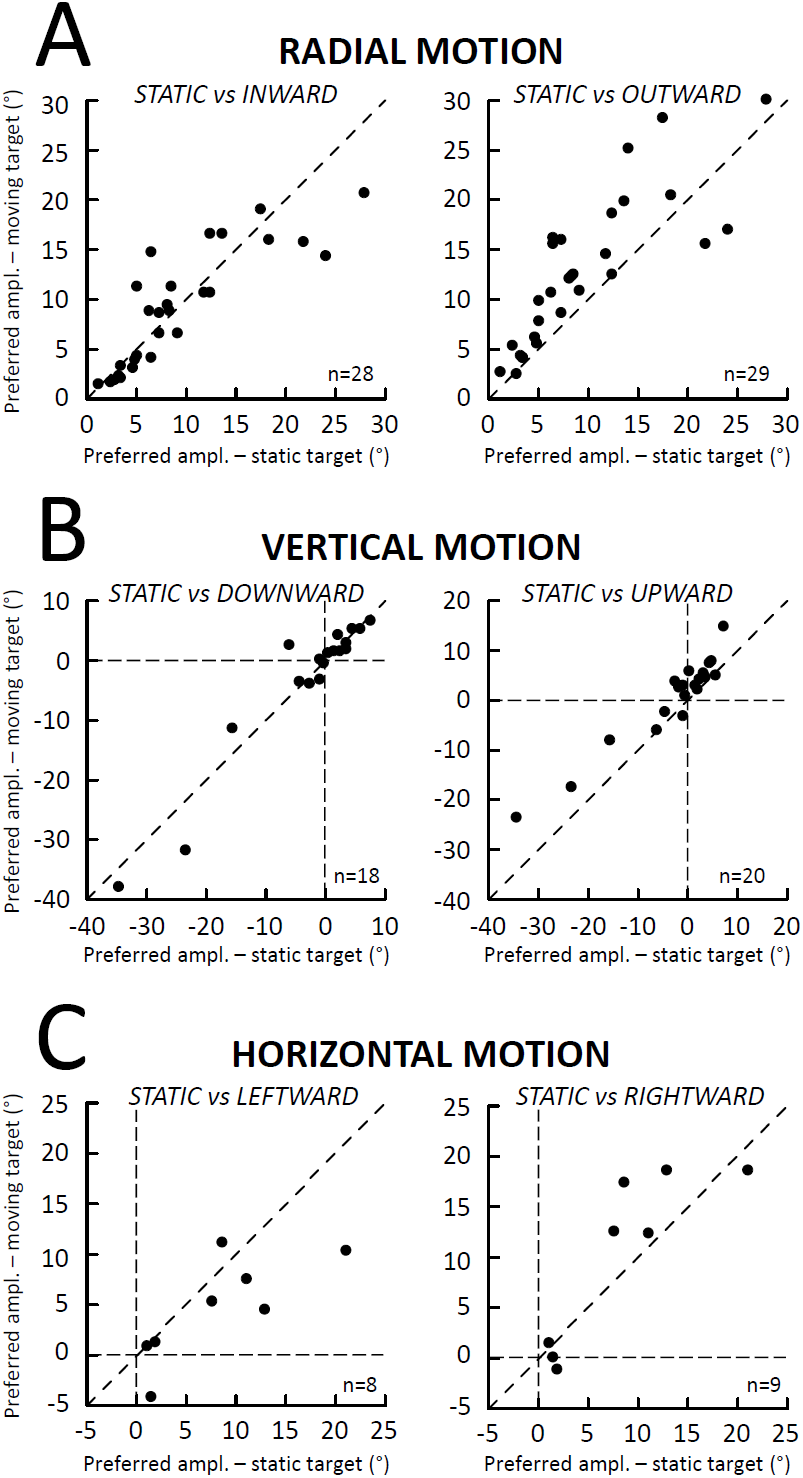
Comparison of the MF center between saccades toward a static target (abscissa) and saccades toward a target (ordinate) moving along the radial axis (A), the vertical axis (B) and the horizontal axis passing through the MF center (C). In B, the MF center could not be estimated for two neurons because of the absence of sharp peak in the curve fitting the relation between firing rate and saccade amplitude.

Finally, when the average firing rates were compared between saccades toward a static and moving target, significant reductions were consistently observed during radial motions (Fig. 8A; −96+/-96 and −101+/-85 spikes/second for inward and outward saccades, corresponding to 24 and 25 % reductions), during vertical motions (Fig. 9B; −54+/-85 and −62+/-110 spikes/second for downward and upward saccades; 15 and 17 % reductions) and during horizontal motions (Fig. 8C; −121+/-98 and −153+/-106 spikes/second for leftward and rightward saccades; 31 and 39 % reductions). Contrary to the suggestion made by Berthoz et al. (1986), the firing rate of SC cells during saccades made toward a moving target is not related to their velocity. Figure 9 shows two examples of cells where the largest difference in MF was found between inward and outward saccades. For the first neuron, when one considers the saccades of amplitudes less than 5 degrees, the firing rate was higher during inward saccades than during outward saccades whereas for saccades of amplitudes larger than 5 degrees, the firing rate was lower during inward saccades than during outward saccades (Fig. 9A; left panel). Yet, the relation between the amplitude and the peak velocity of saccades does not show any difference between the two groups of saccades (Fig. 9A; right panel). For the other neuron, the firing rate was always smaller during outward saccades than during inward saccades (Fig. 9B; left panel) and again, no difference in velocity was observed between the two saccade types (Fig. 9B; right panel). Our results contrast the qualitative impression illustrated in the work of Keller et al. (1996) (see their Figure 1). Perhaps the attenuation reflected as a “shoulder” or double peaks in the velocity waveform was due to an accompanying gaze-evoked blink (Gandhi, 2012).

**Figure 8:**
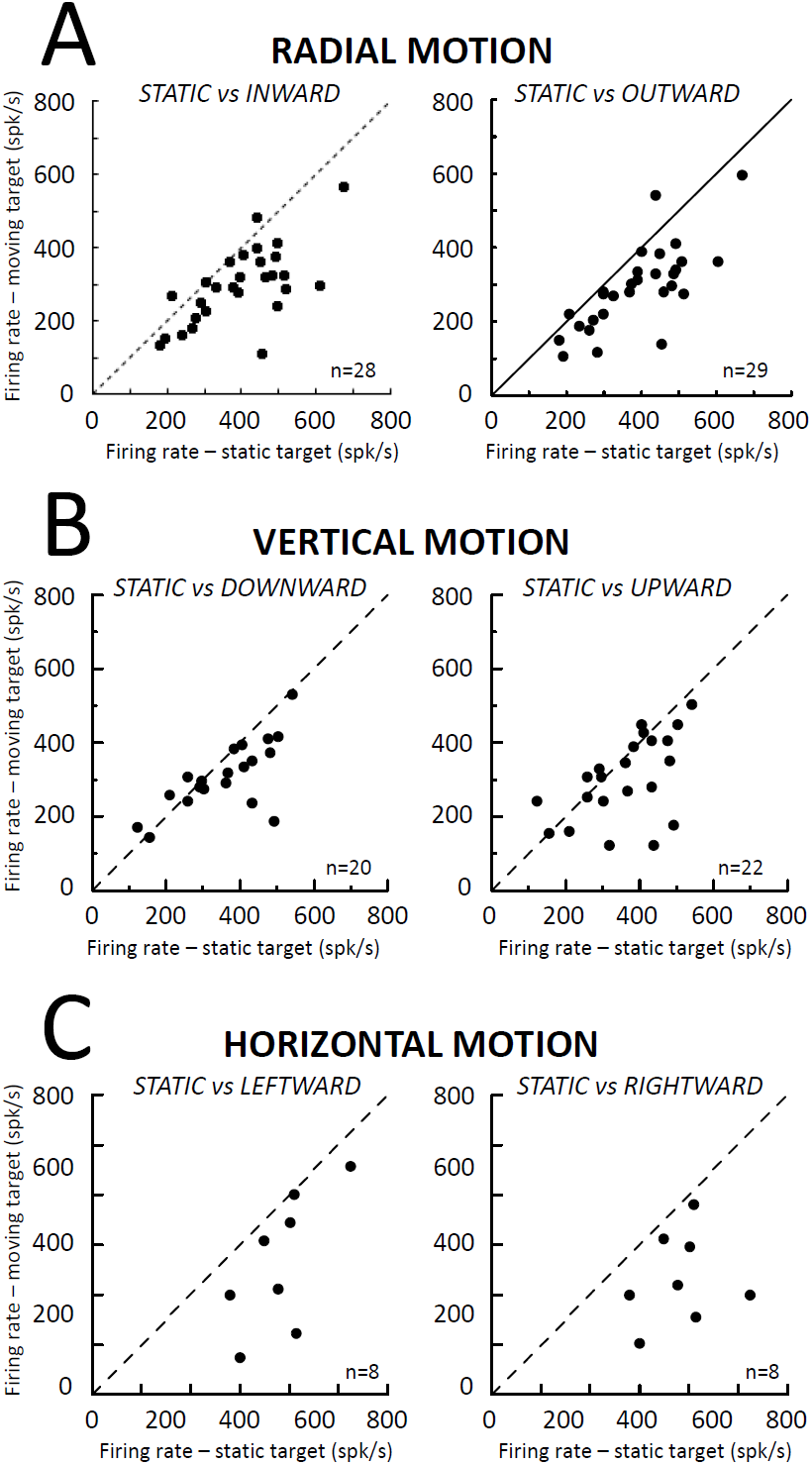
Comparison of the average firing rate (at MF center) of the motor burst between saccades toward a static target (abscissa) and saccades toward a target (ordinate) moving along the radial axis (A), the vertical axis (B) and the horizontal axis passing through the MF center (C).

**Figure 9:**
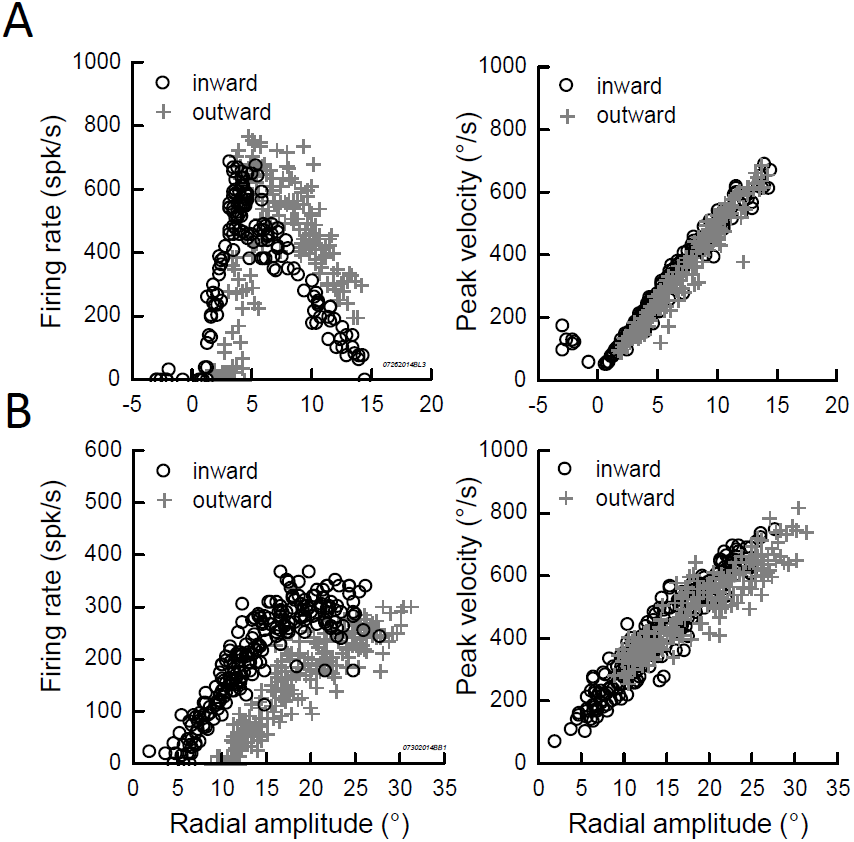
The firing rate of SC cells is not related to the velocity of interceptive saccades. Two examples of cells are shown where the largest difference in MF was found between inward and outward saccades. For the neuron shown in A, the firing rate was higher during inward saccades than during outward saccades of amplitude < 5 degrees, but lower during inward saccades than during outward saccades of amplitude > 5 degrees (left panel). The relation between the amplitude and the peak velocity of saccades does not show any difference between the two groups of saccades (right panel). For the neuron shown in B, the firing rate was lower during outward saccades than during inward saccades (left panel). Again, the relation between the amplitude and the peak velocity of saccades does not show any difference between the two groups of saccades (right panel).

## DISCUSSION

In this work, we studied the movement field (MF) of saccade-related neurons in the SC while monkeys made saccades toward a static or moving visual target. For some neurons, significant shifts were found in the center of the MFs, in their boundaries and in the firing rate. The changes in boundaries and centers indicate that for a given saccade, the population of bursting neurons is not identical between the two types of saccade. However, the shifts were not always observed and their size varied across the cells. When present, they were always in the direction of motion, never in the opposite direction. The absence of shift of boundaries in the direction opposite to the target motion indicates that the SC activity does not issue commands related to upcoming locations of the moving target (no predictive coding). A reduction in the discharge was also observed during interceptive saccades. Unrelated to any change in saccade velocity, this lower firing rate is likely due to the fact that less photons bombarded the retinal cells (and their subsequent recipient visual neurons) when their response field was smoothly "traveled" by a moving target than when it was excited by a static stimulus.

### No predictive coding in the SC for the generation of interceptive saccades

The idea has diffused that the SC would identify the position and speed of an object and, in a predictive and anticipatory manner, trigger the movement required to orient the gaze toward its future location (Berthoz, 2012; Optican & Pretegiani, 2017). Target motion would be “used to predict the future target position so as to assure a spatial lead of the gaze at the saccade end, instead of attempting a precise capture of the target” (Klam et al., 2001). The present study and other works (Hafed et al., 2008; Fleuriet & Goffart, 2012; Quinet & Goffart, 2015a) do not support this suggestion. During the saccade-related burst, the active population does not include cells which code for saccades toward future locations of a moving target. During inward motions, when the target moved from a location outside the MF toward its inside, none of our neurons emitted action potentials that would promote the reduction of saccade amplitude; the outer boundary of their MF did not shift toward larger values of saccade amplitude (Fig. 2B-C and 3B-C). Likewise, during outward motions, when the target moved from a location inside the MF toward a location outside, instead of pausing and facilitating the amplitude increase, the neuron continued to fire, biasing the vector encoded by the population of active neurons toward past locations of the target (Fig. 2B-C and 3B-C) and not to its upcoming locations. In summary, contrary to what would be expected if the SC neurons fired in a predictive manner, the boundaries did not shift in the direction opposite to the target motion. The neurons did not “predictively” fire during saccades toward a target which was going to enter inside their response field. Moreover, their firing persisted when the target, after crossing the response field, moved away from it.

It may be argued that our testing conditions did not favor the possibility of predictive responses because our subjects were not trained to pursue the target, or because the target motion direction and the trials with static and moving targets were pseudo-randomly interleaved. Anticipatory saccades would have likely been observed if the target always moved from the same starting location and in the same direction. Such saccades might even be triggered before the target appears, associated with bursting activities in the SC. However, these premature saccades do not necessarily involve a shift of MF in the direction opposite to the target motion. If the SC activity steers the interceptive saccades like saccades toward a static target, viz., toward the target location (here and now), then the movement fields should overlap between saccades toward static and moving targets.

### The “dual drive” and "remapping" hypotheses

Consistent with the study of Keller et al. (1996), we found that, on average, the MF center shifted in the direction of the target motion during outward saccades (Fig. 7A, rightmost graph). But the shift was small and not consistently observed across all neurons (see examples in Fig. 3C and Fig. 5), comparable to observations made in the frontal eye fields (Cassanello et al., 2008). Should we consider that the generation of outward saccades involves two sub-groups within the active population, with one sub-group composed of neurons which exhibit a shift and another of neurons which do not? This option would require that we consider sub-groups of neurons also for the generation of inward saccades, and likewise for upward and downward saccades. Indeed, the MF center of our example neuron was shifted during inward (Fig. 3B) and upward ones (Fig. 2B) but not during outward (Fig. 3C) or downward saccades (Fig. 2C). The current knowledge of the SC physiology does not support such a segregation (Hall and Moschovakis, 2003; May, 2006; Gandhi and Katnani, 2011). The only known segregation takes place in the pontomedullary and mesencephalic reticular formations, at the level of the premotor neurons which are targeted by the saccade-related SC neurons and which are respectively involved in the generation of the horizontal and vertical components of saccades (Moschovakis et al., 1996; Barton et al., 2003). Therefore, instead of segregation, we propose a continuum of commands within the SC.

Neurophysiological studies indicate that the generation of saccades is under the influence of activity originating in the SC and the CFN. According to the dual drive hypothesis, the MF changes observed during interceptive saccades result from the fact that the saccade-related premotor neurons in the reticular formation are summing commands from the CFN and the SC. Several data are consistent with independent influences of CFN and SC onto the reticular formation, viz., that the fastigial-induced changes in premotor activity do not influence the distribution of active neurons in the SC (see discussion of Quinet and Goffart, 2015b). However, several other observations indicate that the CFN influence on the premotor neurons is modulatory rather than additive (Goffart et al., 2004; Quinet and Goffart, 2007). If the CFN provided a command which compensates for motions of the target away from the vertical meridian (like in Quinet & Goffart, 2015a), one should expect that this supplementary command be constant (or zero) when the target is static. This inference is not consistent with the amplitude-dependent horizontal deviation (ipsipulsion) of vertical saccades during unilateral inactivation of CFN with muscimol (Iwamoto and Yoshida, 2002; Goffart et al., 2004; Quinet and Goffart, 2007). Finally, the dual drive hypothesis considers that the SC encodes the location of the target appearance, overlooking the possibility of subsequent changes in the distribution of active neurons in the SC. However, this view is neither supported by our results nor by the demonstration that the population of active neurons can change during saccades made toward a target which jumps toward a new location (McPeek et al., 2003; Port and Wurtz, 2003).

The shift of the MF boundaries indicates that the locus of activity in the SC is different between identical saccades made toward a static and moving target. The fact that on average the shift is in the same direction as the target motion indicates that the population of active neurons includes commands for generating a saccade toward a past location of the target. The larger shifts of MF centers observed by Keller et al. (1996) are consistent with this view since in their work, the target moved 2 to 3 times faster than in our study. Moreover, the examination of the shift for each individual neuron shows a continuum of neurons ranging from cells which exhibited a shift to cells with no change or very a small shift. Therefore, instead of considering that all SC neurons provide a discrete “snapshot” command and that another drive is added downstream, we propose that the shifts illustrate the fact that the population of active neurons does not change in the SC as fast as the target does in the visual field. Thus, the population in the SC would consist of a continuum of neurons issuing commands, ranging from commands related to antecedent target locations to commands related to its current location. More generally, our study and two others (Hafed et al., 2008; Goffart et al. 2012) show that the SC activity steers the oculomotor system for target foveation, regardless of whether the target is located in the peripheral or central visual field, static or moving. Downstream adjustments for improving the accuracy of foveation are still possible, from the CFN, but from other regions also, since the CFN seems to essentially control their horizontal component (Sato & Noda, 1991; Goffart et al., 2004; Guerrasio et al., 2010; Quinet and Goffart, 2015b). These adjustments would be modulatory and contribute to the spatial and temporal coordination of eye movements with the motion of a visual target in the external world, in a kind of spatial synchronization (Bourrelly et al., 2016).

## REFERENCES

Barton EJ, Nelson JS, Gandhi NJ, Sparks DL (2003) Effects of partial lidocaine inactivation of the paramedian pontine reticular formation on saccades of macaques. J Neurophysiol 90: 372–386.

Berthoz, A. (2012). Simplexity: Simplifying principles for a complex world. G. Weiss, Trans.

Berthoz, A., Grantyn A, Droulez J (1986) Some collicular efferent neurons code saccadic eye velocity. Neurosc Lett 72: 289 294.

Bourrelly C, Quinet J, Cavanagh P, Goffart L (2016) Learning the trajectory of a moving visual target and evolution of its tracking in the monkey. J Neurophysiol 116: 2739–2751.

Bryant CL, Gandhi NJ (2005) Real-time data acquisition and control system for the measurement of motor and neural data. J Neurosci Methods 142: 193–200.

Cassanello CR, Nihalani AT, Ferrera VP (2008) Neuronal responses to moving targets in monkey frontal eye fields. J Neurophysiol 100:1544 –1556.

Fleuriet J, Goffart L (2012) Saccadic interception of a moving visual target after a spatiotemporal perturbation. J Neurosci 32:452– 461.

Fleuriet J, Hugues S, Perrinet L, Goffart L (2011) Saccadic foveation of a moving visual target in the rhesus monkey. J Neurophysiol 105:883– 895.

Fuchs AF, Robinson FR, Straube A (1994) Participation of the caudal fastigial nucleus in smooth-pursuit eye movements. I. Neuronal activity. J Neurophysiol 72: 2714–28.

Gandhi NJ (2012) Interactions between gaze-evoked blinks and gaze shifts in monkeys. Exp Brain Res 216: 321–339.

Gandhi NJ, Katnani HA (2011) Motor functions of the superior colliculus. Annu Rev Neurosci 34: 205–231.

Goffart L, Chen LL, Sparks DL (2004) Deficits in saccades and fixation during muscimol inactivation of the caudal fastigial nucleus in the rhesus monkey. J Neurophysiol 92: 3351–3367, 2004.

Goffart (2017) Saccadic eye movements: Basic neural processes. In Reference Module in Neuroscience and Biobehavioral Psychology, Elsevier.

Guerrasio L, Quinet J, Büttner U & Goffart L (2010) The fastigial oculomotor region and the control of foveation during fixation. J Neurophysiol 103: 1988–2001.

Hafed ZM, Goffart L & Krauzlis RJ (2008) Superior colliculus inactivation causes stable offsets in eye position during tracking. J Neurosci 28: 8124–8137.

Hall WC, Moschovakis AK (2003) The superior colliculus: new approaches for studying sensorimotor integration. CRC Press.

Iwamoto Y, Yoshida K (2002) Saccadic dysmetria following inactivation of the primate fastigial oculomotor region. Neurosci Lett 325: 211–215.

Keller E, Gandhi N, Weir P (1996) Discharge of superior collicular neurons during saccades made to moving targets. J Neurophysiol 76: 3573–3577.

Klam F, Petit J, Grantyn A, Berthoz A (2001) Predictive elements in ocular interception and tracking of a moving target by untrained cats. Exp Brain Res 139: 233–247.

May PJ (2006) The mammalian superior colliculus: laminar structure and connections. Prog Brain Res 151: 321–78.

McPeek RM, Han JH, Keller EL (2003) Competition between saccade goals in the superior colliculus produces saccade curvature. J Neurophysiol 89: 2577–2590.

Moschovakis AK, Scudder CA, Highstein SM (1996) The microscopic anatomy and physiology of the mammalian saccadic system. Prog Neurobiol 50, 133–254.

Optican LM (2009) Oculomotor system: models. In: Encyclopedia of neuroscience (Squire LR, ed.), pp. 25–34. Oxford: Academic.

Optican LM, Pretegiani E (2017) What stops a saccade? Philos Trans R Soc Lond B Biol Sci 372:1718.

Port NL, Wurtz RH (2003) Sequential activity of simultaneously recorded neurons in the superior colliculus during curved saccades. J Neurophysiol 90: 1887–1903.

Quinet J, Goffart L (2007) Head-unrestrained gaze shifts after muscimol injection in the caudal fastigial nucleus of the monkey. J Neurophysiol 98: 3269–3283.

Quinet J, Goffart L (2015a) Does the brain extrapolate the position of a transient moving target? J Neurosci 35: 11780–11790.

Quinet J, Goffart L (2015b) Cerebellar control of saccade dynamics: contribution of the fastigial oculomotor region. J Neurophysiol 113:3323–3336.

Robinson DA (1963) A method of measuring eye movement using a scleral coil in a magnetic field. IEEE Trans Biomed Elect 10: 137–145.

Robinson FR, Fuchs AF (2001) The role of the cerebellum in voluntary eye movements. Annu Rev Neurosci 24: 981–1004.

Sato H, Noda H (1991) Divergent axon collaterals from fastigial oculomotor region to mediodiencephalic junction and paramedian pontine reticular formation in macaques. Neurosci Res 11: 41–44.

Scudder CA, Kaneko CRS, Fuchs AF (2002) The brainstem burst generator for saccadic eye movements: a modern synthesis. Exp. Brain Res 142, 439–462.

Sparks DL (2002) The brainstem control of saccadic eye movements. Nat Rev Neurosci 3: 952–964.

Sparks DL, Lee C, Rohrer WC (1990) Population coding of the direction, amplitude and velocity of saccadic eye movements by neurons in the superior colliculus. Cold Spring Harbor Symp Quant Biol 55:805– 811.

